# Socio-demographic gradients in traumatic brain injury disability, severity and nutrition: a two-level analysis of the Global Burden of Disease Study 2023 and NHANES 2011–2014

**DOI:** 10.64898/2026.10.28.762412

**Authors:** Yukun Hu, Xin Jin, Jiachen Cai, Junxiang Wang

## Abstract

Traumatic brain injury (TBI) is among the most consequential non-fatal outcomes of trauma, yet how its burden, severity and nutritional context are patterned by socio-demographic development remains incompletely characterised. We addressed this question at two levels. Using the Global Burden of Disease (GBD) Study 2023, we quantified the socio-demographic gradients of TBI disability across the five socio-demographic index (SDI) quintiles and characterised the severity composition of the TBI disability burden, its 1990-2023 trend and its cause-specific structure. Using pooled NHANES 2011-2014, we then examined, at the individual level, how reported TBI history among US adults aged 40 years and older varied by income, education, health insurance and food security, and whether dietary differences between adults with and without a TBI history persisted after socio-economic adjustment. The age-standardised TBI disability rate was almost twice as high in the high SDI quintile as in the low SDI quintile (113.68 versus 57.55 per 100,000 in 2023), a gap essentially unchanged since 1990. Moderate or severe injuries accounted for 72.5% of the TBI disability burden in the high SDI quintile versus 84.0-84.6% in the other four quintiles, rising in every quintile between 1990 and 2023. Among US adults, reported TBI history rose with educational attainment (11.9% below high school versus 17.3% above) and was most common in very food-insecure households (23.4%; adjusted odds ratio 1.71, 95% confidence interval 1.39-2.09); income showed no unadjusted gradient but a negative association after mutual adjustment. Unadjusted dietary differences between adults with and without a TBI history largely attenuated after socio-economic adjustment, with only fibre density remaining marginally lower (-0.35 g/1,000 kcal, 95% confidence interval -0.64 to -0.06). The moderate or severe share of the disability burden is a tractable, mortality-free monitoring indicator for injury prevention, and nutritional risk among food-insecure TBI survivors warrants clinical attention.

## 1. Introduction

Traumatic brain injury (TBI) is among the most consequential non-fatal outcomes of trauma worldwide. The Global Burden of Disease (GBD) Study 2016 estimated 27.08 million (95% uncertainty interval [UI] 24.30–30.30 million) new cases of TBI and 55.50 million prevalent cases in 2016, with TBI responsible for 8.1 million (95% UI 6.0–10.4 million) years lived with disability (YLDs), corresponding to an age-standardised rate of 111 (82–141) per 100,000 population [1]. It has further been estimated that up to 69 million people experience TBI each year, with the highest burden in low- and middle-income countries (LMICs) [2]. The GBD Study 2023 — the most recent round, covering 204 countries and territories annually from 1990 to 2023 — continues to provide the reference estimates against which the global TBI burden is tracked [3].

Outcomes after TBI are strongly patterned by socio-economic conditions. A meta-epidemiological analysis of 101 studies from 31 LMICs (59,197 patients) reported a pooled TBI mortality of 16.7% (95% CI 13.7–20.3) — three- to four-fold higher than estimates from high-income countries — with within-LMIC mortality associated with median income, poverty and educational attainment [2]. These differences plausibly reflect a broader care-delivery gap, including limited neurosurgical capacity and restricted access to intracranial pressure monitoring in lower-income settings [4]. Non-fatal burden shows a mirror-image pattern: an analysis of GBD 2021 found that regions with a higher socio-demographic index (SDI) had higher incidence, higher age-standardised incidence rates and higher YLD rates for central nervous system injuries including TBI [5]. Why disability from TBI should be more frequent, in age-standardised terms, in wealthier settings is not obvious; disentangling the contributions of injury frequency, injury severity and post-injury survival requires analyses that stratify the burden by level of socio-demographic development.

Severity is a central, and underused, dimension of the TBI burden. In the GBD framework, TBI is modelled as a nature of injury rather than a cause of death, and is classified into minor and moderate/severe categories; because deaths from head trauma are assigned to the underlying external cause, GBD publishes no TBI-specific mortality or years of life lost [6]. Severity-stratified burden analyses have begun to appear: one analysis of GBD 2021 compared mild and moderate-to-severe TBI across countries and SDI quintiles [7], and a GBD 2023 analysis examined severity-specific trends in the Americas [8]. The global case-count severity proportion has also been described — in 2021, 20,837,466 incident head injuries were reported worldwide, of which 56.63% were moderate/severe [6]. However, to our knowledge, no study has characterised the severity *composition* of the global TBI YLD burden across SDI quintiles — the share of the burden attributable to moderate/severe versus minor injuries — together with its long-term trend and its cause-specific structure. Because this share is derivable from routinely collected estimates, is interpretable without mortality data, and reflects the balance between high-energy and low-energy injury mechanisms, it is a candidate indicator for monitoring whether injury prevention is succeeding in reducing the severity of the injuries being sustained.

Population-level gradients, however, cannot by themselves identify the mechanisms that generate them, and two of the most policy-relevant candidate mechanisms — socio-economic position and nutrition — are only observable in individual-level data. In the United States, the National Health and Nutrition

Examination Survey (NHANES) offers a rare combination of a TBI history item (self-reported head injury with loss of consciousness), detailed socio-economic measures and two days of dietary recall in a nationally representative sample. A previous NHANES analysis compared dietary nutrient intakes between adults with and without a reported head injury with loss of consciousness and found energy-dense diets lower in protein, antioxidants and omega-3 fatty acids among the former [9]; the same item has been used to estimate a weighted prevalence of head injury with loss of consciousness of 15.3% among US adults [10]. To our knowledge, however, no study has asked how reported TBI history itself varies by socio-economic position, whether the previously reported dietary differences persist after adjustment for socio-economic and demographic composition, or how diet quality among TBI survivors varies by food security — the individual-level counterpart of the economic gradients visible in aggregate estimates.

The present study therefore combined two levels of evidence. Using GBD 2023 estimates for the five SDI quintiles and the Global aggregate, we sought to: (i) quantify the between-quintile gradient in the age-standardised TBI YLD rate in 2023 and its change since 1990; (ii) characterise the severity composition of the TBI YLD burden across quintiles and over time; and (iii) describe the external-cause structure and cause-specific severity structure that underlie these gradients. Using pooled NHANES 2011–2014, we additionally sought to: (iv) quantify the economic gradient in reported TBI history among US adults; and (v) compare energy-adjusted nutrient intakes between adults with and without a reported TBI history, before and after socio-economic adjustment, and by food-security status among those with a TBI history.

## Materials and Methods

### 2.1 Study design and data sources

This was a two-level secondary analysis, reported in accordance with the STROBE guideline for observational studies and the GATHER statement for health estimates. The completed STROBE checklist is provided as supplementary file S8 and the GATHER checklist as S9.

**Level 1 (population level).** We re-analysed the Global Burden of Disease Study 2023 (GBD 2023), which estimated the burden of 375 diseases and injuries and of 88 modifiable risk factors for 204 countries and territories, including 660 subnational locations, annually from 1990 to 2023, using DisMod-MR 2.1 for non-fatal outcomes and CODEm for causes of death, with 95% uncertainty intervals (UIs) derived from the 2.5th and 97.5th percentiles of 250 draws [3]. All estimates were retrieved from the GBD Results Tool (https://vizhub.healthdata.org/gbd-results/) on 25 October 2026; the analysis dataset comprised the GBD 2023 results release [11].

Because the unit of analysis in this level was socio-demographic development level rather than individual countries, we stratified all estimates by the five GBD socio-demographic index (SDI) quintiles — a composite indicator of lag-distributed income per capita, mean years of schooling among individuals aged 15 years and older, and the total fertility rate among women younger than 25 years — together with the Global aggregate. National-level estimates were not analysed.

**Level 2 (individual level).** We analysed pooled public-use data from the National Health and Nutrition Examination Survey (NHANES) for the 2011–2012 and 2013–2014 cycles [12], the cycles in which the Taste & Smell questionnaire item on head injury with loss of consciousness (CSQ240) was administered.

NHANES is a nationally representative, multi-stage probability sample of the non-institutionalised US population, combining household interviews with standardised physical examinations and 24-hour dietary recalls in mobile examination centres (MECs).

### 2.2 Case definition and classification

In the GBD framework, traumatic brain injury (TBI) is modelled as a **nature of injury** rather than a cause of death. Consequently, TBI is not included in the GBD cause-of-death hierarchy, and GBD does not publish TBI-specific mortality or years of life lost (YLLs) estimates; deaths from head trauma are assigned to the underlying external cause (e.g., falls, road injuries) [6]. We therefore analysed non-fatal burden and injury severity.

TBI corresponded to the GBD “Head Injuries” nature-of-injury category, encompassing ICD-10 codes F07.2, F07.8, F07.81, F07.89, F07.9, S06, S07, T90.2 and T90.5. The category is subdivided into **minor TBI** (ICD-10 S06.1) and **moderate/severe TBI** (all remaining codes) [6]. These severity classes describe the clinical severity of the injuring event itself and are not measures of treatment outcome.

At the individual level, TBI history was defined by a positive response to the NHANES Taste & Smell questionnaire item CSQ240, “Have you ever had a loss of consciousness because of a head injury?” (yes/no). This item was administered only to participants aged 40 years and older and only in the 2011– 2012 and 2013–2014 cycles; it is self-reported and refers to lifetime history rather than to clinically adjudicated injury.

### 2.3 Outcome measures

**Level 1.** Outcomes were years lived with disability (YLDs), obtained in two forms: (i) age-standardised YLD rates per 100,000 population (age-standardised to the GBD world population standard), used for all between-quintile comparisons; and (ii) absolute YLD counts, used for descriptive burden and for a sensitivity analysis of the severity composition. We did not analyse incidence or prevalence in the GBD dataset, as these data were not extracted for the present analysis.

The primary derived indicator was the **moderate/severe TBI share**, defined as the ratio of the moderate/severe TBI YLD rate to the total Head Injuries YLD rate (age-standardised). This ratio ranges from 0 to 1 and summarises the severity composition of the disability burden of TBI within each SDI quintile and year.

**Level 2.** The outcome for the economic-gradient analysis was reported TBI history (CSQ240 = yes). Exposures of interest were: the family income-to-poverty ratio (INDFMPIR, analysed as quartiles and as a continuous z-score); educational attainment (below high school / high school or equivalent / above high school); health insurance coverage (HIQ011); and adult household food security (FSDAD: full, marginal, low, very low; low and very low combined as “food insecure”). Nutritional outcomes were derived from the day-1 24-hour dietary recall (DR1TOT, supplemented by DR1IFF for individual fatty acids) and expressed as densities per 1,000 kcal (protein, fibre, saturated fatty acids, total sugars, sodium) or per 1,000 kcal for eicosapentaenoic plus docosahexaenoic acid (EPA+DHA; DR1IP205 + DR1IP226, in mg), together with total energy intake (kcal/day) and usual alcohol intake (ALQ130, average drinks per day over the past 12 months; participants who were not past-12-month drinkers or skipped the item were coded 0).

### 2.4 Data extraction

**Level 1.** Using the “Injuries by nature” query context of the GBD Results Tool, we downloaded all estimates for the three nature-of-injury entries “Head Injuries”, “Minor TBI” and “Moderate/Severe TBI”, for the YLDs measure, with metrics “Rate” and “Number”, for both sexes, for Global and the five SDI quintiles, for every year from 1990 to 2023. Rates were downloaded for the age-standardised age group (primary basis for all between-quintile comparisons) and for all ages (sensitivity analyses); counts were downloaded for all ages. Underlying external causes (18 leaf causes: falls; drowning; fire, heat and hot substances; poisonings; exposure to mechanical forces; adverse effects of medical treatment; animal contact; foreign body; electrocution; environmental heat and cold exposure; exposure to forces of nature; other unintentional injuries; road injuries; other transport injuries; self-harm; interpersonal violence; conflict and terrorism; police conflict and executions) were downloaded as a disaggregation of Head Injuries YLDs. The final dataset comprised 10 CSV files (35,088 rows) after excluding non-matching rows.

**Level 2.** We downloaded the public-use XPT files for DEMO (demographics, weights, design variables), CSQ (Taste & Smell), FSQ (food security), HIQ (health insurance), DR1TOT and DR1IFF (day-1 dietary recall and individual fatty acids), SMQ, ALQ, BMX, DIQ, BPQ, DPQ and PAQ (smoking, alcohol, body measures, diabetes, blood pressure, depression screener and physical activity) for both cycles (30 files). The analytic set for the economic-gradient analysis comprised all participants with a valid CSQ240 response (set A, n = 7,399); the dietary analysis additionally required a valid day-1 recall with non-missing energy intake (set D, n = 6,375; 846 with a TBI history).

### 2.5 Statistical analysis

**Level 1.** All GBD analyses were performed in Python 3.13 (standard library only); the analysis script and derived tables are archived with the study materials to ensure reproducibility.

#### Total Head Injuries burden

The 18 leaf causes constitute a complete, non-overlapping partition of Head Injuries; the total Head Injuries YLD rate (and count) was therefore computed as the sum across the 18 leaf causes. UIs for sums were propagated by quadrature assuming independence of cause-specific estimates.

#### Quality control

Before analysis, we verified the internal consistency of the extracted hierarchy by comparing, for each SDI quintile and the Global aggregate at 1990, 2010 and 2023 (18 checks), the sum of the 18 leaf causes for “Head Injuries” with the corresponding sums for “Minor TBI” and “Moderate/Severe TBI” (whose total must equal the Head Injuries total). All checks passed, with a maximum relative discrepancy below 0.001%, confirming correct extraction and additivity of the severity partition.

#### Moderate/severe TBI share

For each quintile and year, the share was computed as the ratio of moderate/severe TBI to total Head Injuries YLD rates. UIs for ratios were derived on the log scale using the delta method; because the numerator and denominator share estimation uncertainty (the Head Injuries total includes the moderate/severe component), the propagated UIs are deliberately conservative and wider than independent-sample intervals. Point estimates are therefore reported as primary, with a count-based (all-ages) ratio as sensitivity analysis.

#### Trends

The estimated annual percentage change (EAPC) in each indicator was calculated for 1990–2023 by least-squares regression of the natural logarithm of the indicator on calendar year: EAPC = 100 × (exp[β] − 1), where β is the regression coefficient; 95% confidence intervals were derived from the standard error of β. EAPC was computed for the total Head Injuries YLD rate and for the moderate/severe TBI share in each quintile.

#### Between-quintile comparisons

Economic gradients were assessed by direct comparison of point estimates across the ordered SDI quintiles (Global as reference), by the absolute and relative differences between the highest and lowest quintiles in 1990 and 2023, and by the change in these differences over time (a decrease in the absolute difference was interpreted as convergence and an increase as divergence). Given the aggregate (ecological) nature of SDI-stratified GBD estimates, no individual-level inference was made, and between-quintile differences are described without formal hypothesis testing.

**Level 2.** NHANES analyses were performed in Python 3.13 using pandas, numpy and pyreadstat, with survey design implemented in the analysis script.

#### Weights and variance

Two sets of sampling weights were used, each divided by two for the pooled four-year analysis: the MEC two-year weight (WTMEC2YR) for analyses not requiring dietary data (set A) and the day-1 dietary weight (WTDRD1) for analyses using the day-1 recall (set D). Variances were estimated by Taylor-series linearisation using the masked design variables (SDMVSTRA, SDMVPSU), with stratum-by-PSU cluster keys that carry the cycle prefix so that identical stratum and PSU numbers in the two cycles are not conflated; the pooled design comprised 29 strata and 61 PSUs. Because the CSQ240 item and the dietary recall were both administered in the MEC, MEC-based weights were used throughout, and participants without MEC dietary data are excluded from set D by design. The variance implementation was validated against a PSU-within-stratum resampling bootstrap (200 replicates for weighted means, 100 for regression coefficients); Taylor-linearisation and bootstrap standard errors agreed to within Monte Carlo error for all four validation estimates (maximum ratio 1.12).

#### Economic gradient

The weighted prevalence of reported TBI history was estimated within levels of each economic factor. Survey-weighted logistic regression was used to estimate adjusted odds ratios (ORs) for each factor separately (adjusted for age, sex and race/ethnicity) and in a mutually adjusted model including all four economic factors simultaneously plus age, sex and race/ethnicity.

#### Nutritional comparisons

Differences in nutrient densities between participants with and without a TBI history were estimated by survey-weighted linear regression, unadjusted and adjusted for age, sex, race/ethnicity, income-to-poverty ratio, education, BMI, smoking, usual alcohol intake, diabetes, hypertension, PHQ-9 score and recreational physical activity. Among participants with a TBI history, nutrient densities were summarised by food-security level, and adjusted associations with the income-to-poverty ratio were estimated; these within-TBI analyses were pre-specified as descriptive and hypothesis-generating given the available sample size.

#### Missing data

Complete-case analysis was used throughout. The PHQ-9 score required all ten items; usual alcohol intake treated non-response as non-drinking. Numbers analysed are reported for each table.

### 2.6 Ethics

The GBD analysis used fully aggregated, publicly available estimates without any individual-level data; no ethics approval was required. The NHANES analysis used public-use, de-identified data files; the NHANES protocol was approved by the National Center for Health Statistics Research Ethics Review Board, and all participants provided written informed consent. No additional ethics approval was required for this secondary analysis.

### 2.7 Use of artificial intelligence

During the preparation of this manuscript, the authors used an artificial intelligence language assistant for language editing and structural drafting. After using this tool, the authors reviewed and edited the content as needed and take full responsibility for the published content. No artificial intelligence tool was used to analyse data or generate results.

## 3. Results

### 3.1 Overall burden of TBI by SDI quintile, 2023

In 2023, TBI (Head Injuries) accounted for 8,624,799 YLDs globally (95% UI 6,828,928–10,420,669), an age-standardised rate of 98.02 per 100,000 (95% UI 77.71–118.33). Burden was strongly graded by socio-demographic development (Table 1, Figure 2). The age-standardised YLD rate was highest in the high-middle SDI quintile (116.73; 95% UI 91.84–141.62) and the high SDI quintile (113.68; 95% UI 88.48–138.89), intermediate in the middle SDI quintile (86.63; 95% UI 70.07–103.19) and low-middle SDI quintile (75.78; 95% UI 61.50–90.05), and lowest in the low SDI quintile (57.55; 95% UI 47.73– 67.37). In absolute terms, the high SDI quintile contributed 3,937,963 YLDs (95% UI 3,051,032– 4,824,893; 45.7% of the global total), followed by the high-middle SDI quintile (2,266,180; 26.3%), whereas the low SDI quintile contributed 745,980 YLDs (95% UI 622,273–869,688; 8.6%).

**Figure 1.**
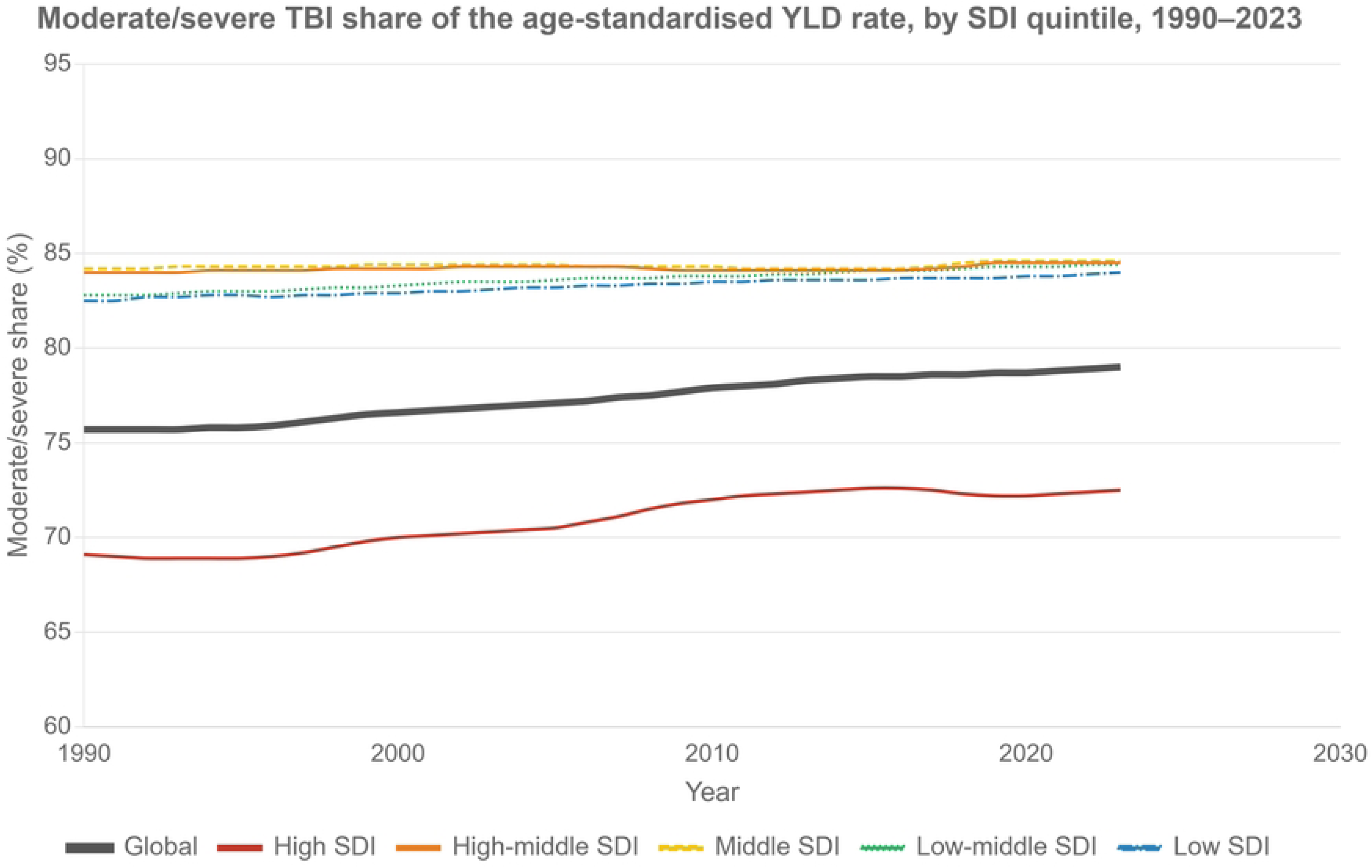
Moderate/severe TBI share (%) of the age-standardised Head Injuries YLD rate, by SDI quintile, 1990–2023. The high SDI quintile runs consistently below the other four quintiles; the share rose in all quintiles over the period. Data: out/tbi_final_table2_severity_share.csv (panel “share” in out/tbi_final_report.html).

**Figure 2.**
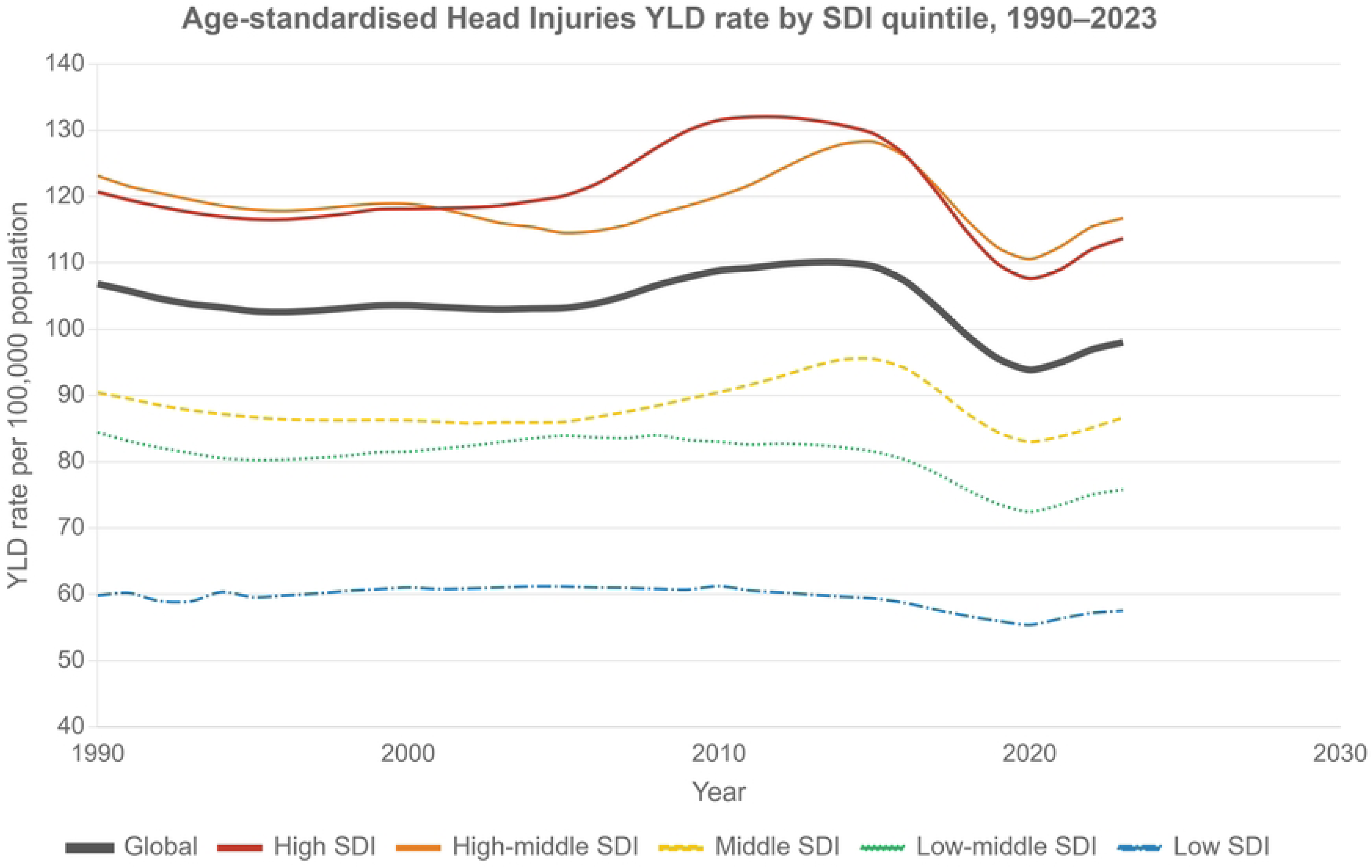
Age-standardised Head Injuries YLD rate per 100,000 population, by SDI quintile, 1990–2023. The high-SDI burden is roughly twice the low-SDI burden throughout, with a stable high-to-low ratio (2.02 in 1990; 1.98 in 2023). Data: analysis/out/tbi_r2_head_yld_rate.csv (panel “head” in out/tbi_final_report.html).

**Table 1.** Burden of TBI (Head Injuries) by SDI quintile, 2023.

| **Location** | **Age-standardised YLD rate per 100,000 (95% UI)** | **Moderate/severe share of YLD rate, %** | **Minor share of YLD rate, %** | **YLD count, all ages (95% UI)** | **Moderate/severe YLD count** |
|---|---|---|---|---|---|
| Global | 98.02 (77.71–118.33) | 79.0 | 21.0 | 8,624,799 (6,828,928–10,420,669) | 6,830,045 |
| High SDI | 113.68 (88.48–138.89) | 72.5 | 27.5 | 3,937,963 (3,051,032–4,824,893) | 2,868,244 |
| High-middle SDI | 116.73 (91.84–141.62) | 84.5 | 15.5 | 2,266,180 | 1,925,847 |
| Middle SDI | 86.63 (70.07–103.19) | 84.6 | 15.4 | 843,857 | 714,886 |
| Low-middle SDI | 75.78 (61.50–90.05) | 84.4 | 15.6 | 823,723 | 693,861 |
| Low SDI | 57.55 (47.73–67.37) | 84.0 | 16.0 | 745,980 | 621,346 |

*Shares are computed from age-standardised YLD rates; counts are all-ages. UI, uncertainty interval; SDI, socio-demographic index; YLD, years lived with disability.*

### 3.2 Severity structure of TBI across SDI quintiles

The moderate/severe TBI share showed a marked socio-demographic gradient (Table 1, Figure 1). In 2023, moderate/severe TBI accounted for 72.5% of the total TBI YLD rate in the high SDI quintile, compared with 84.5% (high-middle), 84.6% (middle), 84.4% (low-middle) and 84.0% (low) in each of the other four quintiles — an absolute difference of approximately 12 percentage points between the high SDI quintile and the remaining quintiles. The global share was 79.0%. The complementary minor TBI share was correspondingly highest in the high SDI quintile (27.5% versus 15.4–16.0% in the other quintiles). The count-based (all-ages) sensitivity analysis reproduced the same ordering and magnitude of the gradient.

Between 1990 and 2023, the moderate/severe share increased in every quintile (Table 2, Figure 1): by 3.3 percentage points globally (75.7% to 79.0%), by 3.4 points in the high SDI quintile (69.1% to 72.5%), and by 0.4–1.6 points in the other four quintiles. EAPCs for the moderate/severe share were positive in all quintiles and largest in the high SDI quintile (+0.193% per year; 95% CI +0.172 to +0.214), followed by the low-middle (+0.061%) and low (+0.053%) SDI quintiles; the global EAPC was +0.148% per year (95% CI +0.142 to +0.154) (Table 3).

**Table 2.** Moderate/severe TBI share (%) of the age-standardised YLD rate, 1990–2023 (selected years)

| **Location** | **1990** | **2000** | **2010** | **2023** | **Change 1990–2023, percentage points** |
|---|---|---|---|---|---|
| Global | 75.7 | 76.6 | 77.9 | 79.0 | +3.3 |
| High SDI | 69.1 | 70.0 | 72.0 | 72.5 | +3.4 |
| High-middle SDI | 84.0 | 84.2 | 84.1 | 84.5 | +0.5 |
| Middle SDI | 84.2 | 84.4 | 84.3 | 84.6 | +0.4 |
| Low-middle SDI | 82.8 | 83.3 | 83.8 | 84.4 | +1.6 |
| Low SDI | 82.5 | 82.9 | 83.5 | 84.0 | +1.5 |

*Changes are computed from the displayed one-decimal values; exact unrounded changes are +0.6, +0.3, +1.6 and +1.5 points for the high-middle, middle, low-middle and low quintiles respectively. The complete annual series (1990–2023) is provided in the supplementary materials.*

**Table 3.** Estimated annual percentage change (EAPC) of TBI indicators, 1990–2023.

| **Location** | **Head Injuries YLD rate EAPC, % per year (95% CI)** | **Moderate/severe share EAPC, % per year (95% CI)** |
|---|---|---|
| Global | −0.127 (−0.27 to +0.01) | +0.148 (+0.142 to +0.154) |
| High SDI | −0.007 (−0.20 to +0.19) | +0.193 (+0.172 to +0.214) |
| High-middle SDI | −0.042 (−0.17 to +0.08) | +0.012 (+0.007 to +0.018) |
| Middle SDI | +0.038 (−0.09 to +0.17) | +0.007 (+0.003 to +0.012) |
| Low-middle SDI | −0.255 (−0.37 to −0.14) | +0.061 (+0.059 to +0.063) |
| Low SDI | −0.169 (−0.25 to −0.09) | +0.053 (+0.050 to +0.055) |

*CI, confidence interval; EAPC, estimated annual percentage change. CIs reflect the regression on GBD point estimates only and do not propagate GBD estimation uncertainty.*

### 3.3 Temporal trends in total TBI disability, 1990–2023

The age-standardised Head Injuries YLD rate declined modestly in most quintiles between 1990 and 2023 (Figure 2): from 106.84 to 98.02 per 100,000 globally (−8.3%), from 120.71 to 113.68 in the high SDI quintile (−5.8%), from 123.15 to 116.73 in the high-middle SDI quintile (−5.2%), from 90.44 to 86.63 in the middle SDI quintile (−4.2%), from 84.43 to 75.78 in the low-middle SDI quintile (−10.2%), and from 59.83 to 57.55 in the low SDI quintile (−3.8%).

EAPCs for the total YLD rate were small and, in the Global aggregate and three of the five quintiles (high, high-middle and middle SDI), not distinguishable from zero (Table 3): −0.127% per year globally (95% CI −0.27 to +0.01), −0.007% (high SDI; 95% CI −0.20 to +0.19), −0.042% (high-middle), +0.038% (middle; 95% CI −0.09 to +0.17), −0.255% (low-middle; 95% CI −0.37 to −0.14) and −0.169% (low; 95% CI −0.25 to −0.09). Thus, the most rapid declines occurred in the two lowest SDI quintiles, whereas the high SDI quintile was essentially static.

The absolute difference in YLD rate between the high and low SDI quintiles narrowed slightly over the period (60.88 per 100,000 in 1990 versus 56.13 in 2023), and the high-to-low ratio was stable (2.02 versus 1.98). In other words, the pronounced between-quintile gap in TBI disability was not substantially reduced during the 33-year period.

### 3.4 Cause structure of TBI by SDI quintile

The contribution of external causes to TBI YLDs differed systematically across SDI quintiles in 2023 (Table 4, Figure 3). Falls were the leading cause in all five quintiles but accounted for a progressively larger share with increasing SDI: 59.5% in the high SDI quintile, 57.5% (high-middle), 49.6% (middle), 49.5% (low-middle) and 40.4% (low). Road injuries were the second leading cause in every quintile, contributing 25.2% (middle), 23.4% (low-middle) and 22.3% (low) in the lower three quintiles versus 19.4% in the high SDI quintile. Interpersonal violence showed the reverse gradient, contributing 13.7% of TBI YLDs in the low SDI quintile versus 7.5% in the high SDI quintile. Conflict and terrorism was a material contributor only in the low SDI quintile (8.7%). The cause-specific pattern in 1990 was qualitatively similar (Table 4).

**Figure 3.**
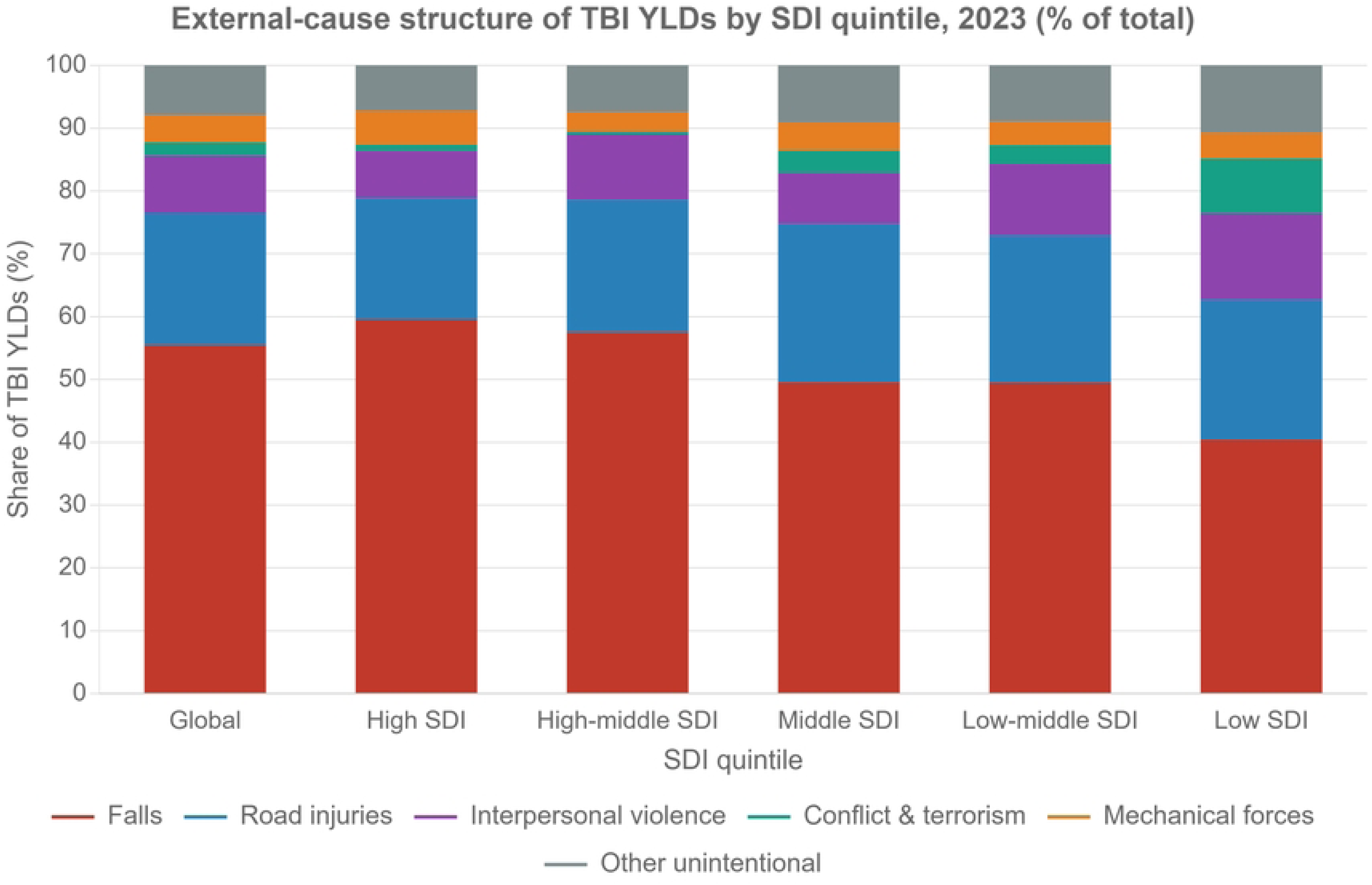
External-cause structure of TBI YLDs (% of total) by SDI quintile, 2023. Falls dominate in high-SDI settings; road injuries, interpersonal violence and conflict contribute progressively more in lower-SDI settings. Data: out/tbi_final_table4_cause_structure.csv (panel “struct” in out/tbi_final_report.html).

**Table 4.** External-cause structure of TBI YLDs by SDI quintile, 2023 (% of total YLD rate)

| **Location** | **Falls** | **Road injuries** | **Interpersonal violence** | **Conflict and terrorism** | **Exposure to mechanical forces** | **Other unintentional injuries** | **Other transport injuries** |
|---|---|---|---|---|---|---|---|
| Global | 55.5 | 21.0 | 9.1 | 2.1 | 4.4 | 4.4 | 1.1 |
| High SDI | 59.5 | 19.4 | 7.5 | 1.0 | 5.4 | 4.1 | 1.2 |
| High-middle SDI | 57.5 | 21.1 | 10.4 | 0.5 | 3.1 | 4.4 | 1.0 |
| Middle SDI | 49.6 | 25.2 | 8.1 | 3.5 | 4.5 | 5.4 | 1.0 |
| Low-middle SDI | 49.5 | 23.4 | 11.4 | 3.0 | 3.7 | 4.4 | 1.2 |
| Low SDI | 40.4 | 22.3 | 13.7 | 8.7 | 4.1 | 5.2 | 1.3 |

*Major causes shown; the remaining leaf causes (drowning; fire, heat and hot substances; poisonings; adverse effects of medical treatment; animal contact; foreign body; electrocution; environmental heat and cold exposure; exposure to forces of nature; self-harm; police conflict and executions) each contributed <2% and are listed in full in the supplementary materials. The 1990 pattern was qualitatively similar.*

### 3.5 Cause-specific severity structure

Among the major causes, the moderate/severe share was consistently lower in the high SDI quintile than in each of the other four quintiles in 2023 (Table 5, Figure 4). For road injuries, the moderate/severe share was 90.4% in the low SDI quintile versus 80.6% in the high SDI quintile; for falls, 82.6% versus 71.1%; for interpersonal violence, 88.9% versus 84.4%. The largest between-quintile contrast was observed for exposure to mechanical forces (75.4% in the low SDI quintile versus 53.3% in the high SDI quintile), although this cause contributed a small fraction of total burden.

**Figure 4.**
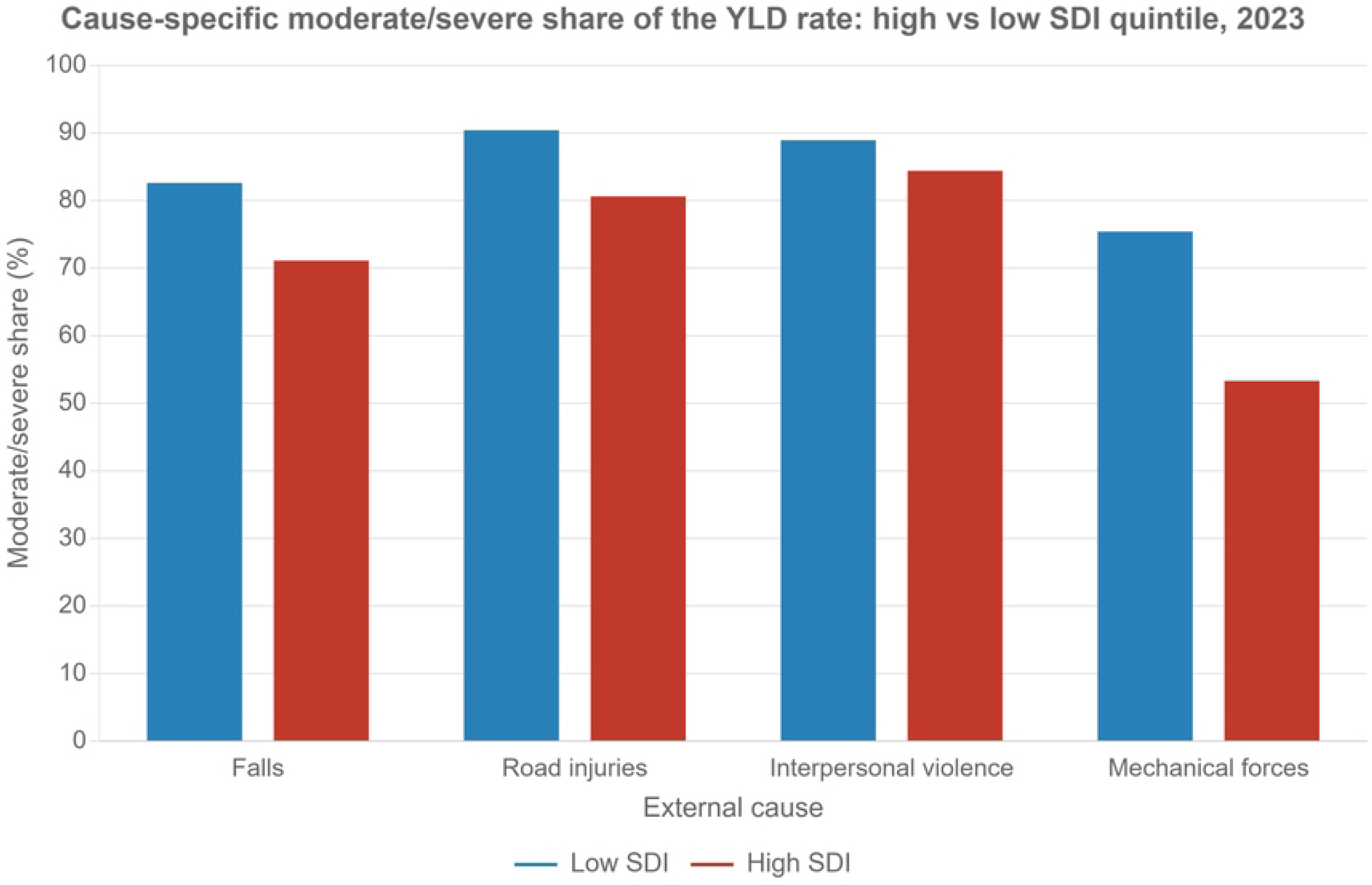
Cause-specific moderate/severe share (%) of the YLD rate, high SDI versus low SDI quintiles, 2023. For every major cause, the share is lower in the high SDI quintile. Data: out/tbi_final_table5_share_by_cause.csv (panel “sbc” in out/tbi_final_report.html).

**Figure 5.**
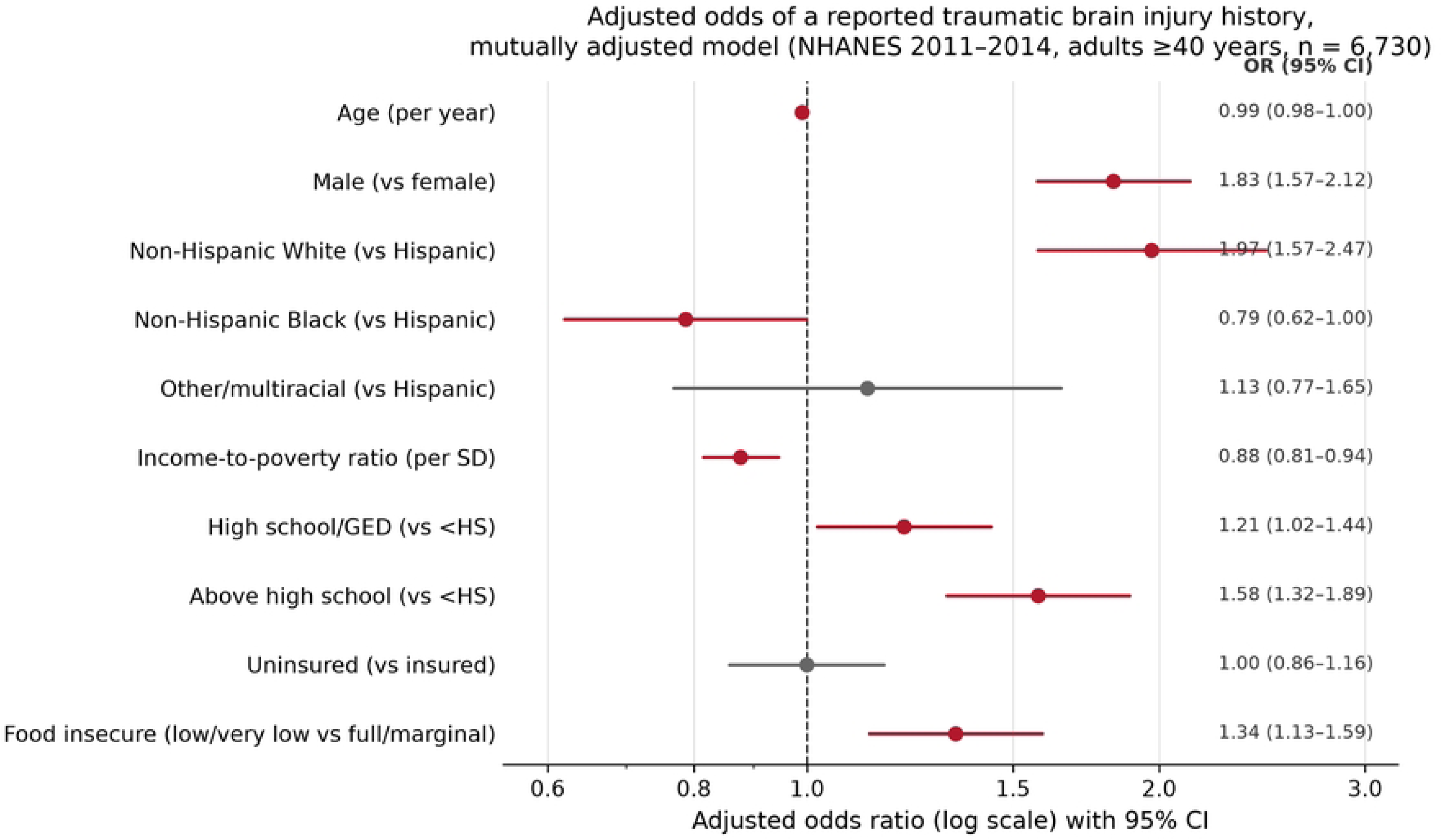
Adjusted odds ratios for a reported TBI history by economic and demographic factors (NHANES 2011–2014). Point estimates with 95% confidence intervals from survey-weighted logistic regression with Taylor-linearisation variances; the income-to-poverty ratio, food security and education terms are from the mutually adjusted model, and demographic terms from the same model. Data: out/tbi_nhanes_tableN2_econ_gradient.csv and out/tbi_nhanes_tableN2b_mutual_model.csv.

**Table 5.** Cause-specific moderate/severe share (%) of the YLD rate by SDI quintile, 2023.

| **Location** | **Falls** | **Road injuries** | **Interpersonal violence** | **Exposure to mechanical forces** |
|---|---|---|---|---|
| Global | 77.0 | 86.6 | 87.7 | 64.4 |
| High SDI | 71.1 | 80.6 | 84.4 | 53.3 |
| High-middle SDI | 82.7 | 91.1 | 89.5 | 77.0 |
| Middle SDI | 82.8 | 90.9 | 89.4 | 76.8 |
| Low-middle SDI | 82.9 | 90.8 | 89.1 | 76.2 |
| Low SDI | 82.6 | 90.4 | 88.9 | 75.4 |

*Uncertainty intervals for cause-specific shares are wide (approximately ±30 percentage points, owing to shared estimation uncertainty between numerator and denominator) and are therefore not displayed; point estimates are reported, and the full results with intervals are in the supplementary materials.*

### 3.6 Economic and nutritional correlates of reported TBI history (NHANES 2011–2014)

The individual-level analysis comprised 7,399 US adults aged 40 years and older with a valid CSQ240 response (948 with a reported TBI history); the weighted prevalence of reported TBI history was 15.7% (95% CI 14.7–16.8). Participants with a TBI history were younger, more often male, more often non-Hispanic White, more highly educated, more often food insecure, and more often current smokers than participants without a TBI history, and had higher PHQ-9 scores (Table 6).

**Table 6.**
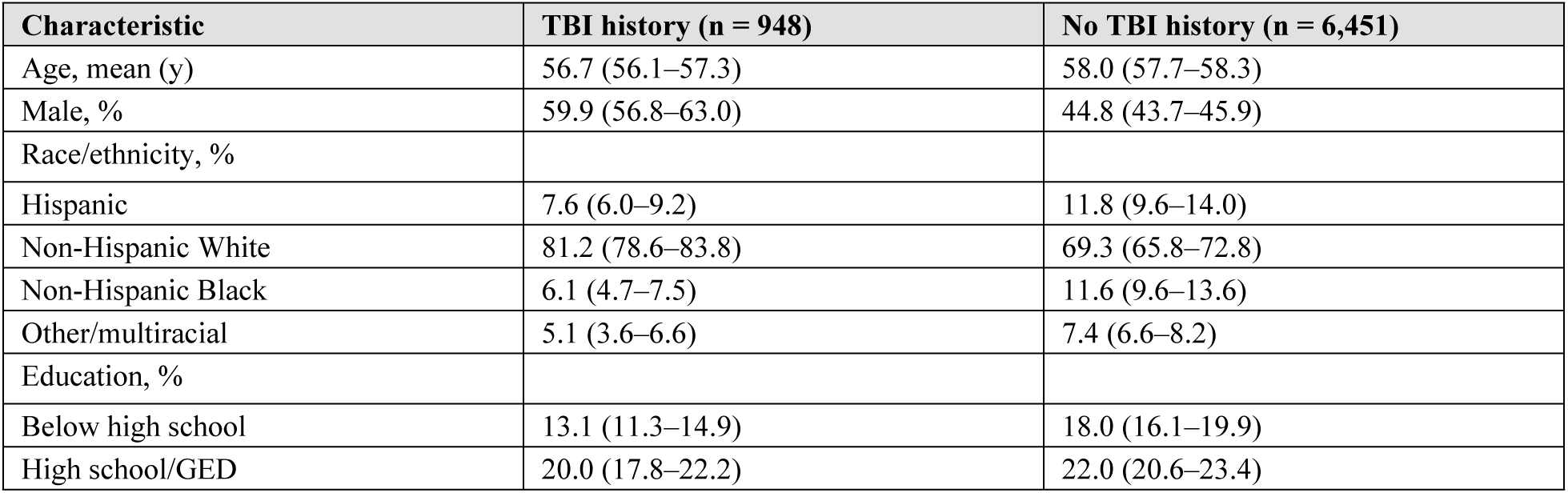
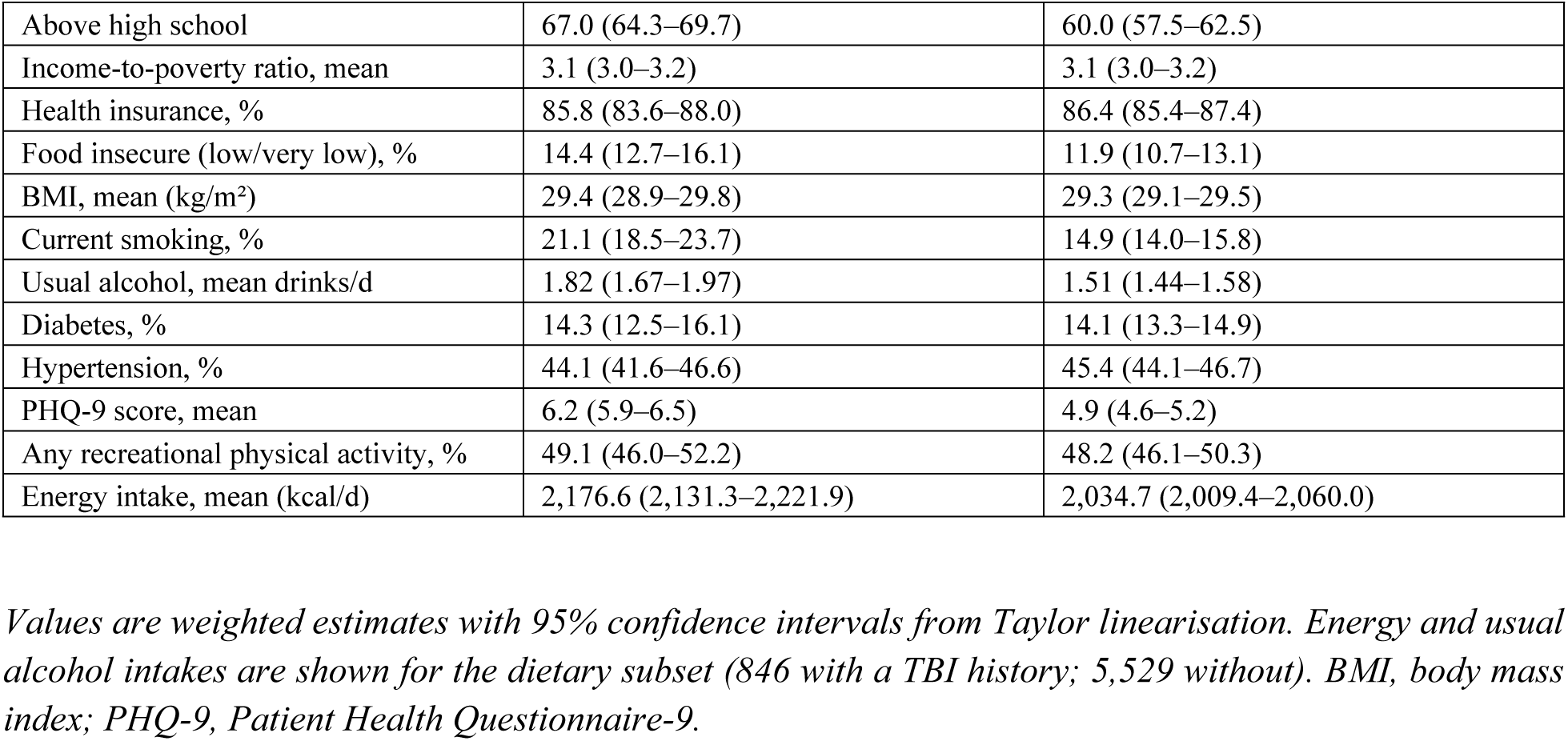
Weighted characteristics of participants with and without a reported TBI history (NHANES 2011–2014, adults ≥40 years)

| **Characteristic** | **TBI history (n = 948)** | **No TBI history (n = 6,451)** |
|---|---|---|
| Age, mean (y) | 56.7 (56.1–57.3) | 58.0 (57.7–58.3) |
| Male, % | 59.9 (56.8–63.0) | 44.8 (43.7–45.9) |
| Race/ethnicity, % |  |  |
| Hispanic | 7.6 (6.0–9.2) | 11.8 (9.6–14.0) |
| Non-Hispanic White | 81.2 (78.6–83.8) | 69.3 (65.8–72.8) |
| Non-Hispanic Black | 6.1 (4.7–7.5) | 11.6 (9.6–13.6) |
| Other/multiracial | 5.1 (3.6–6.6) | 7.4 (6.6–8.2) |
| Education, % |  |  |
| Below high school | 13.1 (11.3–14.9) | 18.0 (16.1–19.9) |
| High school/GED | 20.0 (17.8–22.2) | 22.0 (20.6–23.4) |
| Above high school | 67.0 (64.3–69.7) | 60.0 (57.5–62.5) |
| Income-to-poverty ratio, mean | 3.1 (3.0–3.2) | 3.1 (3.0–3.2) |
| Health insurance, % | 85.8 (83.6–88.0) | 86.4 (85.4–87.4) |
| Food insecure (low/very low), % | 14.4 (12.7–16.1) | 11.9 (10.7–13.1) |
| BMI, mean (kg/m<sup>2</sup>) | 29.4 (28.9–29.8) | 29.3 (29.1–29.5) |
| Current smoking, % | 21.1 (18.5–23.7) | 14.9 (14.0–15.8) |
| Usual alcohol, mean drinks/d | 1.82 (1.67–1.97) | 1.51 (1.44–1.58) |
| Diabetes, % | 14.3 (12.5–16.1) | 14.1 (13.3–14.9) |
| Hypertension, % | 44.1 (41.6–46.6) | 45.4 (44.1–46.7) |
| PHQ-9 score, mean | 6.2 (5.9–6.5) | 4.9 (4.6–5.2) |
| Any recreational physical activity, % | 49.1 (46.0–52.2) | 48.2 (46.1–50.3) |
| Energy intake, mean (kcal/d) | 2,176.6 (2,131.3–2,221.9) | 2,034.7 (2,009.4–2,060.0) |

*Values are weighted estimates with 95% confidence intervals from Taylor linearisation. Energy and usual alcohol intakes are shown for the dietary subset (846 with a TBI history; 5,529 without). BMI, body mass index; PHQ-9, Patient Health Questionnaire-9.*

Reported TBI history was socially patterned (Table 7). Educational attainment showed a positive gradient: prevalence was 11.9% among participants with less than a high-school education, 14.5% among those with a high-school education, and 17.3% among those with more than a high-school education; in adjusted analyses, the odds of a reported TBI history were lower for participants with less than a high-school education (OR 0.62, 95% CI 0.54–0.72) or a high-school education (OR 0.80, 0.71–0.91) than for those with more than a high-school education. Food security showed a non-monotonic pattern: prevalence was 14.9% in fully food-secure households, 18.9% in marginally food-secure households (adjusted OR 1.30, 1.04–1.63), 14.6% in low-food-security households (OR 0.95, 0.78–1.15) and 23.4% in very-low-food-security households (OR 1.71, 1.39–2.09). The income-to-poverty ratio showed no unadjusted gradient (prevalence 15.2–16.6% across quartiles) and health insurance showed no association (adjusted OR 0.92, 0.79–1.08 for uninsured versus insured).

**Table 7.** Economic correlates of reported TBI history among US adults ≥40 years (NHANES 2011– 2014)

| **Factor and level** | **n** | **TBI history prevalence, % (95% CI)** | **Adjusted OR (95% CI)\*** |
|---|---|---|---|
| Income-to-poverty ratio quartile |  |  |  |
| Q1 (lowest) | 1,174 | 16.5 (14.4–18.6) | 1.04 (0.86–1.26) |
| Q2 | 1,593 | 15.2 (13.6–16.8) | 0.94 (0.81–1.08) |
| Q3 | 1,850 | 16.0 (14.0–18.0) | 0.99 (0.83–1.19) |
| Q4 (highest) | 2,117 | 16.6 (15.5–17.7) | Reference |
| Education |  |  |  |
| Below high school | 1,921 | 11.9 (10.3–13.5) | 0.62 (0.54–0.72) |
| High school/GED | 1,652 | 14.5 (13.1–15.9) | 0.80 (0.71–0.91) |
| Above high school | 3,819 | 17.3 (16.1–18.5) | Reference |
| Health insurance |  |  |  |
| Insured | 6,149 | 15.7 (14.9–16.5) | Reference |
| Uninsured | 1,250 | 16.3 (14.1–18.5) | 0.92 (0.79–1.08) |
| Food security |  |  |  |
| Full | 5,361 | 14.9 (14.0–15.8) | Reference |
| Marginal | 754 | 18.9 (16.1–21.7) | 1.30 (1.04–1.63) |
| Low | 724 | 14.6 (12.3–16.9) | 0.95 (0.78–1.15) |
| Very low | 500 | 23.4 (19.7–27.1) | 1.71 (1.39–2.09) |

*Adjusted for age, sex and race/ethnicity. OR, odds ratio.*

#### Adjusted for age, sex and race/ethnicity. OR, odds ratio

In a mutually adjusted model including all four economic factors (n = 6,730), the income-to-poverty ratio showed a negative association with reported TBI history (OR 0.877 per standard deviation, 95% CI 0.815–0.945) once educational attainment and food insecurity were taken into account, whereas food insecurity (OR 1.34, 1.13–1.59) and above-high-school education (OR 1.58, 1.32–1.89 relative to below high school) remained positively associated; health insurance remained unassociated (OR 1.00, 0.86– 1.16). Male participants had higher odds than female participants (OR 1.83, 1.57–2.13), non-Hispanic White participants had higher odds than Hispanic participants (OR 1.97, 1.57–2.47), and odds declined slightly with age (OR 0.990 per year, 0.984–0.996).

In the dietary subset (n = 6,375; 846 with a TBI history), participants with a TBI history had higher unadjusted energy intake (+141.9 kcal/day, 95% CI +92.2 to +191.6), higher total sugar density (+2.44 g/1,000 kcal, +0.31 to +4.56), lower fibre density (−0.45 g/1,000 kcal, −0.72 to −0.18), lower EPA+DHA density (−13.7 mg/1,000 kcal, −22.7 to −4.6) and higher usual alcohol intake (+0.31 drinks/day, +0.19 to +0.43) than participants without a TBI history (Table 8). After adjustment for age, sex, race/ethnicity, income-to-poverty ratio, education, BMI, smoking, usual alcohol intake, diabetes, hypertension, PHQ-9 score and recreational physical activity, the differences in energy, total sugars, EPA+DHA and alcohol attenuated to the null (energy −10.8, 95% CI −84.5 to +62.9; total sugars +2.22, −0.64 to +5.07; EPA+DHA −5.9, −18.1 to +6.3; alcohol −0.02, −0.17 to +0.13), whereas fibre density remained marginally lower among participants with a TBI history (−0.35 g/1,000 kcal, −0.64 to −0.06); no other nutrient differed significantly between the two groups after adjustment.

**Table 8.** Nutrient intakes of participants with and without a reported TBI history (dietary subset, NHANES 2011–2014)

| **Nutrient** | **TBI history (n = 846)** | **No TBI history (n = 5,529)** | **Unadjusted difference (95% CI)** | **Adjusted difference (95% CI)\*** |
|---|---|---|---|---|
| Protein (g/1,000 kcal) | 39.13 (38.31–39.95) | 39.71 (39.27–40.15) | −0.58 (−1.34 to +0.19) | +0.15 (−0.84 to +1.15) |
| Fibre (g/1,000 kcal) | 8.52 (8.20–8.84) | 8.97 (8.80–9.14) | −0.45 (−0.72 to −0.18) | −0.35 (−0.64 to −0.06) |
| Saturated fat (g/1,000 kcal) | 12.23 (11.91–12.55) | 11.95 (11.83–12.07) | +0.28 (−0.08 to +0.64) | +0.24 (−0.18 to +0.66) |
| Total sugars (g/1,000 kcal) | 54.53 (52.51–56.55) | 52.09 (51.30–52.88) | +2.44 (+0.31 to +4.56) | +2.22 (−0.64 to +5.07) |
| Sodium (mg/1,000 kcal) | 1,660.2 (1,625.5–1,694.9) | 1,698.9 (1,684.3–1,713.5) | −38.7 (−79.8 to +2.4) | −13.8 (−57.6 to +30.0) |
| EPA+DHA (mg/1,000 kcal) | 40.67 (32.08–49.26) | 54.34 (49.91–58.77) | −13.7 (−22.7 to −4.6) | −5.9 (−18.1 to +6.3) |
| Usual alcohol (drinks/d) | 1.82 (1.67–1.97) | 1.51 (1.44–1.58) | +0.31 (+0.19 to +0.43) | −0.02 (−0.17 to +0.13) |
| Energy (kcal/d) | 2,176.6 (2,131.3–2,221.9) | 2,034.7 (2,009.4–2,060.0) | +141.9 (+92.2 to +191.6) | −10.8 (−84.5 to +62.9) |

*Adjusted for age, sex, race/ethnicity, income-to-poverty ratio, education, BMI, smoking, usual alcohol intake, diabetes, hypertension, PHQ-9 score and recreational physical activity; the alcohol row is not adjusted for usual alcohol intake (the outcome itself). CI, confidence interval; EPA+DHA, eicosapentaenoic plus docosahexaenoic acid.*

Among participants with a TBI history and dietary data, nutrient densities showed descriptive gradients by food-security status (Table 9). EPA+DHA density fell from 44.5 mg/1,000 kcal in fully food-secure households to 38.5 in marginally food-secure and 22.9 in low/very-low-food-security households; fibre density followed a similar pattern (8.92, 6.76 and 7.88 g/1,000 kcal), and total sugar density rose (53.2, 60.5 and 57.6 g/1,000 kcal). In adjusted analyses among participants with a TBI history (n = 571 with complete covariates), the income-to-poverty ratio was positively associated with protein density (+0.73 g/1,000 kcal per unit PIR, 95% CI +0.07 to +1.39) and inversely associated with total sugar density (−2.03 g/1,000 kcal, −3.33 to −0.72); no other nutrient density showed a statistically significant socio-economic gradient. Given the modest sample size and the number of nutrients examined, these within-TBI analyses are interpreted as descriptive and hypothesis-generating.

**Table 9.** Nutrient densities by food-security status among participants with a reported TBI history (NHANES 2011–2014, n = 846)

| **Nutrient** | **Full food security (n = 571)** | **Marginal food security (n = 100)** | **Low/very low food security (n = 172)** | **Adjusted β per 1-unit PIR (95% CI)\*** |
|---|---|---|---|---|
| Protein (g/1,000 kcal) | 39.91 (39.03–40.79) | 35.27 (32.23–38.31) | 37.91 (36.82–39.00) | +0.73 (+0.07 to +1.39) |
| Fibre (g/1,000 kcal) | 8.92 (8.53–9.31) | 6.76 (6.16–7.36) | 7.88 (7.39–8.37) | +0.07 (−0.14 to +0.28) |
| Saturated fat (g/1,000 kcal) | 12.06 (11.76–12.36) | 12.70 (11.65–13.75) | 12.78 (12.29–13.27) | −0.01 (−0.18 to +0.17) |
| Total sugars (g/1,000 kcal) | 53.17 (50.91–55.43) | 60.45 (54.10–66.80) | 57.55 (53.91–61.19) | −2.03 (−3.33 to −0.72) |
| Sodium (mg/1,000 kcal) | 1,665.9 (1,623.6–1,708.1) | 1,603.4 (1,545.5–1,661.3) | 1,636.4 (1,568.9–1,704.0) | −17.9 (−45.6 to +9.8) |
| EPA+DHA (mg/1,000 kcal) | 44.49 (33.31–55.67) | 38.48 (24.10–52.86) | 22.86 (17.67–28.05) | +5.0 (−0.4 to +10.4) |
| Usual alcohol (drinks/d) | 1.68 (1.54–1.82) | 2.41 (2.06–2.76) | 1.89 (1.49–2.29) | −0.04 (−0.15 to +0.07) |
| Energy (kcal/d) | 2,174.3 (2,120.3–2,228.3) | 2,230.2 (2,080.6–2,379.8) | 2,118.9 (2,008.8–2,229.0) | +18.4 (−18.8 to +55.6) |

*Adjusted for age, sex, race/ethnicity, education, BMI, smoking, usual alcohol intake (except in the alcohol row), PHQ-9 score and recreational physical activity. CI, confidence interval; PIR, income-to-poverty ratio.*

## 4. Discussion

### 4.1 Principal findings

In the population-level analysis of GBD 2023, we found that the non-fatal burden of TBI, measured as YLDs, was almost twice as high in the high SDI quintile as in the low SDI quintile (113.7 versus 57.5 per 100,000 in 2023), that this gap was essentially unchanged between 1990 and 2023 despite modest absolute declines everywhere, and that the severity composition of the TBI disability burden is systematically graded by socio-demographic development: moderate/severe injuries accounted for 72.5% of the TBI YLD rate in the high SDI quintile versus 84.0–84.6% in the other four quintiles in 2023 — a pattern that, to our knowledge, has not previously been described for the share of the YLD burden across SDI quintiles. The cause structure underlying these gradients was equally systematic: falls dominated in high-SDI settings, whereas road injuries, interpersonal violence and conflict contributed progressively more in lower-SDI settings.

At the individual level, the NHANES 2011–2014 analysis indicated that reported TBI history among US adults aged 40 years and older was socially patterned in three respects: it increased with educational attainment (11.9% below high school versus 17.3% above high school), was more common in very food-insecure households (23.4%; adjusted OR 1.71) and, after mutual adjustment, decreased with the income-to-poverty ratio (OR 0.877 per SD); it showed no association with health insurance. Participants with a TBI history consumed less fibre and omega-3 fatty acids and more energy, sugar and alcohol than participants without one in unadjusted analyses; after adjustment for socio-economic and demographic composition, only the fibre difference persisted (−0.35 g/1,000 kcal, 95% CI −0.64 to −0.06). Among participants with a TBI history, omega-3 density was lowest in food-insecure households (22.9 versus 44.5 mg/1,000 kcal in low/very-low versus fully food-secure households), with adjusted income gradients for protein and total sugar densities in this underpowered subgroup.

### 4.2 Interpretation of the socio-demographic gradients

The finding that TBI disability is *more* frequent, in age-standardised terms, in wealthier settings may appear counter-intuitive, but it is consistent with the structure of the GBD estimation framework and with the wider injury literature. Three mechanisms are plausible and not mutually exclusive.

First, the gradient partly reflects **case ascertainment**. Mild TBI, which is readily missed without imaging and structured follow-up, constituted 27.5% of the TBI YLD rate in the high SDI quintile but only 15.4– 16.0% elsewhere. A substantial share of the high-SDI excess burden is therefore likely to represent injuries that are identified and coded in high-income settings but remain uncounted in lower-income settings — a measurement gradient superimposed on the biological one. The same logic applies to the global rise in the moderate/severe share over the study period, which may in part reflect improved capture of severe cases rather than a true worsening of injury severity; this ambiguity cannot be resolved with GBD data alone.

Second, the gradient reflects the **age distribution of injury risk**. Falls — the leading cause of TBI in every quintile — are strongly concentrated at older ages, and the GBD world population standard assigns substantial weight to these ages. In high-SDI settings, where low-energy falls in older adults dominate the cause structure (59.5% of TBI YLDs, versus 40.4% in the low SDI quintile), the age-specific injury rates at the ages carrying the greatest standard-population weight are plausibly higher than in settings where the burden is concentrated at younger ages, which raises the age-standardised rate. The same demographic structure also inflates absolute counts: the high SDI quintile contributed 45.7% of global TBI YLDs in 2023.

Third, the gradient reflects **differential survival after injury**. Low-SDI settings carry a substantially higher case fatality from TBI — a pooled mortality of 16.7% (95% CI 13.7–20.3) in low- and middle-income countries versus roughly three- to four-fold lower estimates in high-income countries [2] — so that a smaller fraction of injured patients survive to accrue years lived with disability. The YLD gradient documented here is therefore, if anything, a *lower bound* on the true inequality in TBI outcomes: the mortality dimension, which GBD does not publish for TBI as a nature of injury, is almost certainly steeper still.

The severity-structure gradient admits a more specific interpretation. Because the minor/moderate-severe classification describes the injuring event rather than its treatment, the 12-percentage-point excess in the high SDI quintile’s *minor* share most plausibly reflects two components: (i) a **cause-mix effect** — low-SDI TBI is more often produced by high-energy mechanisms (road injuries contributing 22.3–25.2% of YLDs in the lower three quintiles versus 19.4% in the high SDI quintile; interpersonal violence and conflict adding a further 13.7% and 8.7% in the low SDI quintile); and (ii) a **within-cause effect** — even for the same external cause, injuries were more often moderate/severe in lower-SDI quintiles (road injuries 90.4% versus 80.6%; falls 82.6% versus 71.1%), consistent with differences in exposure to protective factors (helmets, restraints, vehicle and workplace safety standards) and in the built environment. We note, however, that severity shares are derived quantities sensitive to the underlying cause distribution and to coding practice, and that the wide uncertainty intervals we report reflect genuine shared estimation uncertainty rather than sampling error alone.

The individual-level analysis provides a suggestive, if partial, correspondence with these mechanisms. The positive gradient in *reported* TBI history by educational attainment is consistent with the case-ascertainment mechanism operating at the level of individual recall and reporting: within a single country and health system, adults with more education reported a head injury with loss of consciousness more often, plausibly because milder or more remote injuries are remembered, recognised and disclosed more readily — the same direction of bias that is hypothesised to inflate measured burden in higher-SDI populations. Conversely, the elevated prevalence in very food-insecure households (23.4%) is consistent with a genuine exposure mechanism: material deprivation is associated with hazardous living and working environments, older housing, and greater exposure to violence. That income showed no unadjusted gradient but a negative association once education and food insecurity were taken into account underscores that these dimensions of socio-economic position are not interchangeable and that single-indicator analyses can misstate the direction of a gradient. The food-security pattern was non-monotonic — elevated in both the marginal and very-low categories but not in the low category — which cautions against a simple dose-response reading and may reflect the heterogeneity of the very-low group and the limited precision of the estimates.

The nutritional findings sharpen the interpretation of the economic gradient. A previous NHANES analysis reported energy-dense diets lower in protein, antioxidants and omega-3 fatty acids among US adults with a self-reported head injury with loss of consciousness [9]. Our analysis reproduces the unadjusted pattern — higher energy intake and lower fibre and omega-3 densities among participants with a TBI history — and shows that it is almost entirely attributable to socio-economic and demographic composition: after adjustment, only fibre density remained marginally lower among participants with a TBI history (−0.35 g/1,000 kcal, 95% CI −0.64 to −0.06). The nutritional disparity associated with TBI in these data is therefore, in the main, a socio-economic disparity — the individual-level counterpart of the population-level SDI gradient, and a reminder that unadjusted comparisons of diet between disease groups can confound the disease with its social determinants. What remains after adjustment is a small fibre difference and a descriptive signal among TBI survivors themselves: omega-3 density was roughly halved in food-insecure households (22.9 versus 44.5 mg/1,000 kcal), a pattern consistent with the broader literature on food insecurity and diet quality, with adjusted income gradients for protein and total sugar densities also emerging in this underpowered subgroup. Omega-3 fatty acids and other antioxidant nutrients are biologically plausible candidates for modulating post-injury neuroinflammation and secondary injury, and the absence of dietary guidance for TBI survivors has previously been noted [9]; our findings identify food-insecure TBI survivors as a group in whom nutritional risk is likely to be concentrated.

The absence of meaningful improvement deserves emphasis. Over 33 years, age-standardised TBI YLD rates fell by 3.8–10.2% across quintiles, with EAPCs indistinguishable from zero in the Global aggregate and in three of five quintiles, and the high-to-low ratio moved only from 2.02 to 1.98. In contrast to several other major causes of health loss, for which GBD 2023 documents double-digit declines in age-standardised rates in recent decades [3], TBI prevention appears to have captured remarkably little of the gains achieved elsewhere in global health. Whether this reflects the difficulty of modifying the distal determinants of injury (road systems, alcohol, violence, conflict, ageing) or a relative neglect of injury prevention in global health governance is beyond what these data can establish, but the finding itself is a signal that merits attention from policy-makers.

### 4.3 Implications

For high-income and ageing societies, the dominant message is that TBI is increasingly a **disease of older adults sustaining low-energy falls** — the leading cause of TBI in every SDI quintile in our data, and one that rises steeply with age — and that prevention effort should shift accordingly: fall-risk assessment, home and institutional environments, anticoagulation management in older patients, and geriatric trauma pathways. For neurosurgeons in such settings, the corollary is a rising caseload of chronic subdural haematoma and other fall-related pathologies in the elderly, with the attendant questions of surgical decision-making in frail patients.

For low- and middle-income settings, the analysis points to a **double burden of mechanism and severity**: a larger share of TBI arises from road traffic, violence and conflict, and those injuries are more often severe. This compounds the documented deficits in neurosurgical and neurocritical-care capacity in lower-income settings — scarce neurosurgical services, limited intracranial pressure monitoring and restricted access to timely surgical decision-making [4] — into a widening absolute gap in survival and disability. Road-safety legislation and enforcement, helmet and restraint programmes, violence-prevention initiatives, and investment in tiered trauma systems remain the highest-yield interventions; our data suggest that such investments would also shift the severity composition of TBI, not merely its frequency.

The severity structure itself may be useful as a **monitoring indicator**. Because it is derivable from routinely collected GBD estimates and is interpretable without mortality data, the moderate/severe share offers health systems a tractable proxy for the trajectory of injury prevention — specifically, whether prevention is reducing the severity of the injuries being sustained — complementing conventional frequency metrics.

At the individual level, the nutritional findings carry a complementary implication for clinical practice: because the dietary differences associated with a TBI history in these data reflect socio-economic composition rather than the injury itself, nutritional assessment and dietary counselling after TBI should be targeted by socio-economic and food-security screening rather than by injury history alone. The concentration of low omega-3 and protein density among food-insecure TBI survivors — if confirmed in adequately powered studies — would make food-insecure survivors a priority group for nutritional intervention and for trials of dietary modification after injury.

### 4.4 Strengths and limitations

This analysis has several strengths. It uses the most recent GBD round (2023), extending the TBI-specific GBD analyses published to date — all based on GBD 2021 [5,6,7] — to the current estimation round; it introduces an SDI-stratified, share-based analysis of the severity composition of the YLD burden, with cause-specific decomposition and 1990–2023 trends, going beyond the severity-stratified burden estimates previously reported by SDI quintile [7] and beyond the region-specific (Americas) severity analysis of GBD 2023 [8]; it rests on a complete, non-overlapping partition of causes with internal-consistency verification (18/18 checks, maximum relative discrepancy <0.001%); and it pairs the population-level analysis with a nationally representative individual-level analysis using pre-specified survey-weighted methods with validated variance estimation.

Several limitations must be considered. First, the GBD design is **ecological**: SDI quintiles are aggregates, and no individual-level inference is possible; the observed gradients describe populations, not persons, and are subject to the ecological fallacy. Second, GBD estimates themselves carry uncertainty, and the EAPC confidence intervals we report reflect only the regression on the point estimates; they do not propagate GBD estimation uncertainty. The log-linearity assumption underlying EAPC may also be violated over 33 years, particularly around the 2020–2021 COVID-19 disruption, which affected data availability and injury patterns worldwide. Third, GBD does not publish TBI-specific mortality or YLLs because TBI is modelled as a nature of injury; our conclusions therefore concern non-fatal burden, and the mortality dimension is discussed only by reference to external evidence. Fourth, the severity classification describes the injuring event, not treatment outcome, and is sensitive to diagnostic and coding practices that differ across settings; the severity gradient we report should be read as a mixture of true differences in injury mechanisms and ascertainment differences. Fifth, the uncertainty intervals for the moderate/severe share are deliberately conservative: because the Head Injuries total includes the moderate/severe component, the numerator and denominator of the ratio are not independent, and propagating their estimation uncertainty as if independent widens the intervals (95% UIs span roughly 40–50 percentage points; for example, 72.5% [95% UI 53.4–98.5] for the high SDI quintile in 2023). Sixth, the count-based sensitivity analysis uses all-ages counts, which are not age-standardised and therefore reflect differences in population size and age structure in addition to risk. Finally, we did not analyse incidence or prevalence in the GBD dataset; the “incidence paradox” (higher incidence alongside higher disability in high-SDI settings) is supported here only indirectly, through previously published GBD 2021 estimates [5].

The NHANES analysis has its own limitations. The TBI history item is **self-reported** and refers to lifetime history of head injury with loss of consciousness rather than to clinically adjudicated injury; recall and disclosure are likely to be imperfect and, as our education gradient suggests, socially patterned, so the prevalence and gradient estimates describe reported rather than true histories. The item was administered **only in 2011–2014 and only to adults aged 40 years and older**, so the findings are not generalisable to younger adults or to more recent periods, and the cross-sectional design precludes any causal or temporal interpretation. The analysis is restricted to the **United States**, and its individual-level gradients should not be read as a test of the population-level SDI gradient. Dietary intakes are self-reported and subject to well-described misreporting, including systematic under-reporting of energy and alcohol; nutrient densities mitigate but do not eliminate this. Complete-case analysis was used throughout, and the within-TBI socio-economic analyses (n = 846; 571 with complete covariates) were powered only for large effects, so their adjusted findings — including the significant protein and sugar gradients — should be treated as hypothesis-generating rather than confirmatory. Finally, the usual-alcohol measure treated non-response as non-drinking, a conservative assumption that may modestly attenuate alcohol estimates.

### 4.5 Conclusion

In 2023, TBI accounted for 8.6 million YLDs worldwide, with an age-standardised burden nearly twice as high in the high SDI quintile as in the low SDI quintile — a gap that has barely moved since 1990. The severity composition of TBI is graded in the opposite direction to frequency: moderate/severe injuries make up a markedly smaller share of the burden in high-SDI settings, tracking the shift from high-energy mechanisms (road traffic, violence, conflict) to low-energy falls. At the individual level, reported TBI history in the United States is patterned by education and food insecurity, and the dietary differences observed among TBI survivors are largely socio-economic in origin, with only fibre density remaining marginally lower after adjustment. These patterns indicate that global TBI prevention has lagged behind other areas of health improvement, that the inequality in TBI outcomes is larger in mortality than in disability, and that both mechanism-specific prevention (road safety, fall prevention, violence reduction) and the integrity of the injury-care chain remain unfinished business across the development spectrum — with nutritional risk among disadvantaged TBI survivors an additional, and modifiable, dimension for clinicians and health systems to address.

## Acknowledgments

The authors thank the Institute for Health Metrics and Evaluation (IHME) for providing the Global Burden of Disease estimates and the National Center for Health Statistics (NCHS) for the National Health and Nutrition Examination Survey public-use data. The funders had no role in the study design, data collection and analysis, decision to publish, or preparation of the manuscript.

## Author Contributions

Yukun Hu: Conceptualization, Methodology, Formal Analysis, Data Curation, Funding Acquisition, Writing - Original Draft. Xin Jin: Data Curation, Validation, Visualization. Jiachen Cai: Writing - Review & Editing. Junxiang Wang: Conceptualization, Supervision, Writing - Review & Editing. All authors reviewed and approved the final manuscript. *(Contribution statement to be confirmed by all authors before submission.)*

## Data Availability

All data underlying the results reported in this study are publicly available. GBD 2023 estimates are available from the Institute for Health Metrics and Evaluation (IHME) Global Burden of Disease Results Tool (https://vizhub.healthdata.org/gbd-results/, GBD 2023 release). NHANES 2011-2014 public-use files are available from the CDC National Center for Health Statistics (https://wwwn.cdc.gov/nchs/nhanes/). The analysis scripts and all derived tables are provided as supplementary file S10 and are also available from the corresponding author upon reasonable request.

## Abbreviations

BMI: body mass index
CDC: Centers for Disease Control and Prevention
CI: confidence interval
CODEm: Cause of Death Ensemble modeling
DisMod-MR: Disease Model – Meta-Regression
EAPC: estimated annual percentage change
EPA+DHA: eicosapentaenoic plus docosahexaenoic acid
GATHER: Guidelines for Accurate and Transparent Health Estimates Reporting
GBD: Global Burden of Disease
GED: General Educational Development
HIC: high-income country
ICD-10: International Classification of Diseases, 10th revision
ICP: intracranial pressure
LMIC: low- and middle-income country
MEC: mobile examination centre
NCHS: National Center for Health Statistics
NHANES: National Health and Nutrition Examination Survey
OR: odds ratio
PHQ-9: Patient Health Questionnaire-9
PIR: income-to-poverty ratio
SDI: socio-demographic index
STROBE: Strengthening the Reporting of Observational Studies in Epidemiology
TBI: traumatic brain injury
UI: uncertainty interval
YLD: years lived with disability
YLL: years of life lost.

## Supporting Information

S1 File. Complete 1990-2023 annual series of the moderate or severe TBI share of the age-standardised YLD rate, by SDI quintile.

S2 File. Full 18-leaf external-cause structure of TBI YLDs, by SDI quintile, 2023, with uncertainty intervals.

S3 File. Full cause-specific moderate or severe share by SDI quintile, 2023, with uncertainty intervals.

S4 File. Mutually adjusted economic-gradient logistic model for reported TBI history (all four economic factors).

S5 File. Bootstrap validation of Taylor-linearisation variance estimation.

S6 File. Quality-control checks of the GBD cause hierarchy (18 of 18 passed).

S7 File. NHANES 2011-2014 public-use file manifest.

S8 File. STROBE checklist for this observational study.

S9 File. GATHER checklist for the health-estimate analysis.

S10 File. Analysis scripts and derived tables (archive).

## STROBE Statement-Checklist of items that should be included in reports of observational studies

**Manuscript:** Socio-demographic gradients in traumatic brain injury disability, severity and nutrition: a two-level analysis of the Global Burden of Disease Study 2023 and NHANES 2011-2014

*Completed checklist. The study has two components: a population-level secondary analysis of aggregated GBD 2023 estimates (reported according to GATHER, supplied separately) and an individual-level cross-sectional secondary analysis of pooled NHANES 2011-2014 public-use microdata (this checklist). Items concerning primary data generation are marked “Not applicable” with justification*.

| # | Item | Reported in |
|---|---|---|
| 1a | Indicate the study's design with a commonly used term in the title or the abstract | Title and Abstract (“two-level analysis”; Level 2 = cross-sectional secondary analysis of pooled NHANES cycles); Methods 2.1. |
| 1b | Provide in the abstract an informative and balanced summary of what was done and what was found | Abstract (structured: Background, Methods, Results, Conclusions; both levels reported). |
| 2 | Explain the scientific background and rationale for the investigation being reported | Introduction (paragraphs 1–4; Level 2 rationale in paragraph 4, citing [9] and [10]). |
| 3 | State specific objectives, including any prespecified hypotheses | Introduction (final paragraph, aims (i)–(v)); study protocol (archived with study materials). |
| 4 | Present key elements of study design early in the paper | Methods 2.1 (two-level secondary analysis; Level 2 = pooled cross-sectional, survey-weighted). |
| 5 | Describe the setting, locations, and relevant dates, including periods of recruitment, exposure, follow-up, and data collection | Methods 2.1 and 2.4 (NHANES 2011–2014, United States, non-institutionalised civilian population; MEC examinations and 24-hour dietary recalls; public-use files accessed 25 October 2026; Table S7). |
| 6a | Cross-sectional study — give the eligibility criteria, and the sources and methods of selection of participants | Methods 2.2–2.4 (CSQ240 administered to participants aged ≥40 years in the 2011–2012 and 2013–2014 cycles; analytic set A n = 7,399 with valid CSQ240; set D n = 6,375 with valid day-1 recall; complete-case analysis). |
| 6b | For matched studies, give matching criteria and number of exposed/unexposed | Not applicable (unmatched cross-sectional analysis). |
| 7 | Clearly define all outcomes, exposures, predictors, potential confounders, and effect modifiers | Methods 2.3 (outcome: reported TBI history, CSQ240; exposures: income-to-poverty ratio, education, health insurance, adult food security; outcomes for dietary analyses: nutrient densities per 1,000 kcal, energy, usual alcohol); 2.5 (confounder sets for each model). |
| 8 | For each variable of interest, give sources of data and details of methods of assessment; describe comparability if more than one group | Methods 2.3–2.4 and Table S7 (variable-level provenance: CSQ240 self-report lifetime history; INDFMPIR, DMDEDUC2, HIQ011, FSDAD; DR1TOT day-1 recall with DR1IFF fatty acids summed by SEQN; ALQ130 usual alcohol with non-response coded 0). |
| 9 | Describe any efforts to address potential sources of bias | Methods 2.5 (survey weights, design-based variance, energy-adjusted nutrient densities to mitigate misreporting, mutual adjustment of economic factors); Discussion 4.4 (self-report, recall, disclosure and misreporting biases discussed with direction). |
| 10 | Explain how the study size was arrived at | Methods 2.4 (all eligible participants in the two cycles were included; within-TBI analyses pre-specified as descriptive given the available sample size, Methods 2.5). |
| 11 | Explain how quantitative variables were handled in the analyses; describe which groupings were chosen and why | Methods 2.3 (income-to-poverty ratio analysed as quartiles and as a z-score; education in three levels; food security in four levels with low/very low combined as food insecure), 2.5 (z-score standardisation for the mutually adjusted model). |
| 12a | Describe all statistical methods, including those used to control for confounding | Methods 2.5 (survey-weighted logistic regression with Taylor-linearisation variances; mutually adjusted model; survey-weighted linear regression for nutrient differences, unadjusted and adjusted for age, sex, race/ethnicity, income-to-poverty ratio, education, BMI, smoking, alcohol, diabetes, hypertension, PHQ-9 and physical activity). |
| 12b | Describe any methods used to examine subgroups and interactions | Methods 2.5 and Results 3.6 (food-security-stratified descriptions among participants with a TBI history; no formal interaction tests — pre-specified as descriptive). |
| 12c | Explain how missing data were addressed | Methods 2.5 (complete-case analysis; PHQ-9 required all ten items; usual alcohol treated non-response as non-drinking; numbers analysed reported for each table). |
| 12d | Cohort study — explain how loss to follow-up was addressed | Not applicable (cross-sectional design; no follow-up). |
| 12e | Describe any sensitivity analyses | Methods 2.5 (variance implementation validated against a PSU-within-stratum resampling bootstrap, Table S5; weighted TBI prevalence compared with published estimates, Discussion 4.4); Level 1 count-based ratio sensitivity (Methods 2.5). |
| 13a | Report numbers of individuals at each stage of the study | Methods 2.4 and Table S6 (DEMO rows 19,931; CSQ respondents ≥40 years 7,418; TBI+ 948; analytic sets A 7,399 and D 6,375); Tables 6–9 report per-table numbers. |
| 13b | Give reasons for non-participation at each stage | Not applicable (secondary analysis of public-use data; item non-response documented in Table S6 and handled by complete-case analysis, Methods 2.5). |
| 13c | Consider use of a flow diagram | Table S6 (numeric flow: respondents, TBI status, analytic sets). |
| 14a | Give characteristics of study participants and information on exposures and potential confounders | Results 3.6 and Table 6 (weighted characteristics by TBI status); Tables 7–9. |
| 14b | Indicate number of participants with missing data for each variable of interest | Methods 2.4–2.5 and Table S6 (complete-case numbers: set A 7,399 of 7,418 CSQ respondents; mutual model 6,730; within-TBI adjusted analyses 571 of 846). |
| 14c | Cohort study — summarise follow-up time | Not applicable (cross-sectional). |
| 15 | Report numbers of outcome events or summary measures of exposure | Results 3.6 (weighted prevalence of reported TBI history 15.7%; prevalences by economic factor, Table 7; nutrient densities, Tables 8–9). |
| 16a | Give unadjusted estimates and, if applicable, confounder-adjusted estimates and their precision; make clear which confounders were adjusted for | Results 3.6, Tables 7–9 (unadjusted and adjusted estimates with 95% CIs; adjustment sets stated in each table footnote and Methods 2.5). |
| 16b | Report category boundaries when continuous variables were categorised | Methods 2.3 (PIR quartiles; education categories; food-security categories); Table 7. |
| 16c | If relevant, consider translating estimates of relative risk into absolute risk | Results 3.6 and Table 7 (prevalences reported by exposure level alongside adjusted ORs, enabling absolute-risk interpretation). |
| 17 | Report other analyses done — subgroups, interactions, sensitivity analyses | Results 3.6 (food-security-stratified dietary descriptions among TBI+ participants); Table S5 (variance validation). |
| 18 | Summarise key results with reference to study objectives | Discussion 4.1 (Principal findings, both levels), mapped to aims (i)–(v). |
| 19 | Discuss limitations of the study, taking into account sources of potential bias or imprecision; discuss direction and magnitude | Discussion 4.4 (NHANES limitations: self-report, 2011–2014 and ≥40-years restriction, cross-sectional design, US-only generalisability, dietary misreporting, complete-case and power limitations — each with direction). |
| 20 | Give a cautious overall interpretation of results considering objectives, limitations, multiplicity of analyses, results from similar studies, and other relevant evidence | Discussion 4.2 (mechanisms framed as plausible, not demonstrated), 4.3, 4.4, 4.5; within-TBI analyses explicitly labelled descriptive and hypothesis-generating. |
| 21 | Discuss the generalisability (external validity) of the study results | Discussion 4.4 (US-only, adults ≥40 years, 2011–2014; individual-level gradients not a test of the population-level SDI gradient); weighted prevalence benchmarked against published estimates [10]. |
| 22 | Give the source of funding and the role of the funders | Title page; Declarations (Funding: SYW2026096; funder had no role). |

